# Biofouling on aquaculture mesh: disentangling seasonal and environmental effects with eDNA

**DOI:** 10.1101/2025.11.20.689612

**Authors:** Tian Tian, Xavier Pochon, Peter Bell, Maren Wellenreuther

## Abstract

Biofouling presents ecological and economic challenges for marine industries, yet patterns of community development in relevant environments remain underexplored. In this study, we examined seasonal (winter vs summer) and spatial (sheltered vs exposed) patterns in eukaryotic biofouling across six coastal sites in New Zealand. Using standardised mesh substrates and environmental DNA (eDNA) metabarcoding with the 18S rRNA gene, we characterised over 10,000 Amplicon Sequence Variants (ASVs) across 216 samples, revealing 541 species from 249 taxonomic classes. Exposed sites exhibited marked seasonal contrasts: winter communities were dominated by hydroids (e.g., *Coryne eximia*), while summer samples shifted to amphipod-rich assemblages (e.g., *Jassa slatteryi*). Sheltered sites showed more stable biomass and diversity but greater site-level variability, especially in summer. Temperature and wind fetch emerged as key environmental drivers, with larger seasonal fluctuations observed under high-exposure conditions. Notably, several dominant species were non-native, underscoring the importance of early detection. These findings highlight the value of molecular monitoring for predicting biofouling risks and informing site-specific antifouling strategies in dynamic marine environments.

## Introduction

Biofouling is a natural process in marine ecosystems but also one of the most significant challenges in marine-related industries, including maritime transport and shipping, aquaculture, and other coastal anthropogenic activities (Flemming et al. 2009; Fitridge et al. 2012). The biofouling succession process usually begins with an initial microbial biofilm which is subsequently followed by the establishment of larger macro-organisms such as algae and barnacles (Callow and Callow 2002). Biofouling communities contribute to habitats, biodiversity, and food webs (Lewis 1998). However, this biomass also creates challenges for industries by increasing weight and surface damage on vessels and aquaculture structures, spreading harmful species, and ultimately leading to higher maintenance costs (Dürr and Thomason 2009). Specifically, the additional costs for global shipping fuel due to hull fouling are estimated to be 9% of the fleet’s fuel consumption per year (Hewitt 2023). In marine aquaculture, approximately 5-10% of the total aquaculture production costs are used for biofouling management (Lane and Willemsen 2004). These challenges indicate the pressing need for efficient biofouling assessment methods to develop both cost-effective antifouling tools as well as enhance general management strategies in the marine environment and its relevant industries.

Over the last years, numerous antifouling tools have been developed and applied to mitigate impacts (Cao et al. 2011). However, many of them remain inefficient and pose potential pollution risks to marine ecosystems (Bannister et al. 2019). For example, some commonly used antifouling coatings contain compounds, such as copper or organotin, that are toxic to many non-target species (Amara et al. 2018; Lagerström et al. 2020). Likewise, physical removal approaches risk releasing harmful species into the surrounding environment (Woods et al. 2012). Modern alternative technologies, such as ultrasound disruption and silicone-based materials, may reduce negative environmental impacts, yet their maintenance costs, scalability and long-term effectiveness are still unclear (Hu et al. 2020; Trickey et al. 2022). Furthermore, standardised and universal management can be challenging due to high biofouling community variability driven by environmental and seasonal conditions (Vinagre et al. 2020). Local regulations, risk levels, and cost constraints vary between industries and regions, indicating site-specific and adaptive approaches are necessary to facilitate effective management.

Fine-scale monitoring of biofouling communities is critical to understand how the changing environments influence their growth and to identify potential problematic species. As such, resulting data can inform community modelling and future prediction, which facilitates targeted and efficient management (Bannister et al. 2019). Data composition analyses can further support the early detection of newly arrived species and provide insights into the key drivers of successional patterns. By tracking the presence and absence of species and growth of key biofouling species, high-risk periods and locations for fouling can be predicted (Wu et al. 2023). This allows for more specific approaches that ultimately deliver effective management actions, while reducing unnecessary treatments and environmental impacts. For example, biofouling monitoring can inform aquaculture site and farming season selection, enable long-term modelling of harmful species distribution, and guide cleaning schedules and antifouling measures (Sylvester et al. 2011; Atalah et al. 2017).

Until recently, marine biofouling monitoring studies have primarily relied on traditional methods such as visual occlusion estimation (Sievers et al. 2014; Wrange et al. 2020). While widely used, these approaches suffer from limited taxonomic resolution and higher costs, particularly for cryptic or closely related species. Over the past two decades, molecular tools, such as next generation sequencing (NGS) and eDNA metabarcoding, have emerged as powerful alternatives for community analysis (Taberlet et al. 2012; Behjati and Tarpey 2013). By using genetic markers to simultaneously identify large taxonomic assemblages from different environments, eDNA metabarcoding offers more accurate and comprehensive insights into community composition (Valentini et al. 2009; Ruppert et al. 2019). Although widely adopted in ecological and environmental assessments of biofouling communities (Pochon, Zaiko, et al. 2015; von Ammon et al. 2018; Pearman et al. 2021), its application to marine biofouling in industrial contexts remains limited, highlighting a significant opportunity for improving monitoring.

In this study, we used eDNA metabarcoding to characterise eukaryotic biofouling communities on standard industrial mesh across two seasons (winter and summer) and two exposure regimes (sheltered and exposed) in New Zealand coastal waters. By combining biomass measurements with molecular analyses, we investigated both seasonal succession and short-term recruitment patterns. Our objectives were to: (1) assess biomass accumulation and alpha diversity across environmental gradients; (2) examine seasonal and spatial shifts in community composition and beta diversity; and (3) evaluate the influence of key environmental drivers, particularly temperature and wind exposure. We hypothesised that biofouling dynamics would vary significantly with season and exposure, and that these insights could inform more effective, site-specific biofouling management strategies for marine industries.

## Materials and Methods

### Experimental design to characterise biofouling dynamics

To investigate seasonal and spatial patterns in biofouling community structure, we selected six sampling sites in the northern region of New Zealand’s South Island. Three exposed sites located in Tasman Bay (Exposed 1-3; E1-E3) and three sheltered sites in Pelorus Sound (Sheltered 1-3; S1-S3) served as replicates for the two exposure conditions (Figure 1A). Exposure in this context was determined by comparing relative differences in wind fetch between these sites, which was calculated by averaging the distances from the sampling site to the shores in all directions (Seers 2018). Sampling was conducted across two seasons, with winter spanning from June to September 2023 and summer from December 2023 to March 2024. Each site was sampled three times per season to capture both seasonal succession and monthly recruitment patterns.

**Figure 1.**
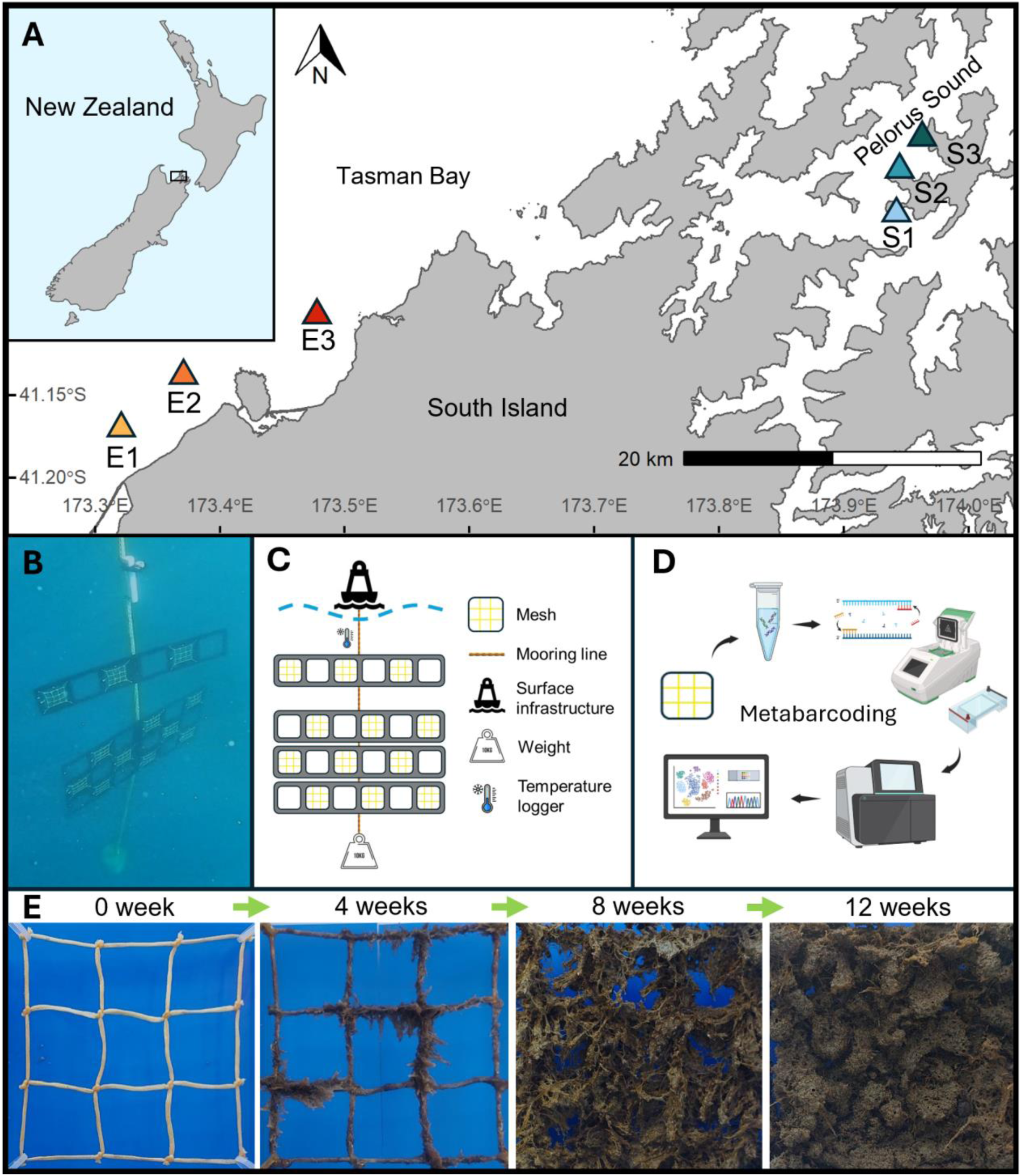
Sampling sites, deployment structures, metabarcoding procedures and successional samples. (A) Locations and sites where sampling conducted: Tasman Bay (exposed sites E1-3) and Pelorus Sound (sheltered sites S1-3); (B) actual underwater image of deployed structures at S1; (C) sample deployment structure design with the 6 × 1 monthly recruitment panel and the 6 × 3 seasonal successional panel (Vecteezy 2025; ClipartMax 2018); (D) metabarcoding procedures (BioRender 2025); (E) seasonal succession samples at the initial deployment and at each sampling (0-12 weeks) at E1 in winter.

Biofouling settlement was assessed using standardised material: 2.5 mm diameter Dyneema® mesh (knotted T90 square mesh, bar length = 33 mm, supplied by Hampidjan New Zealand Ltd), commonly used in marine industries, especially aquaculture. For each treatment, three replicate 15 cm × 15 cm mesh squares were attached to three small plastic inner frames using cable ties and then attached to a 6 × 1 plastic array frame (manufactured by Nelson Plastics Ltd) (Figure 1B & 1C). For each site, four array frames were attached to a mooring line, including one for the monthly recruitment treatment (5 m below the sea surface) and three attached together for the seasonal succession treatment (15 cm below the monthly recruitment frame) (Figure 1B & 1C). The mooring lines were either bottom mounted with surface floats (exposed sites) or attached and hung from surface infrastructure (sheltered sites). One environmental data logger (HOBO^®^ Pendant Mx Temperature Logger, Onset, USA) was deployed under water at each site and set to record every 10 min. Metabarcoding preparations were performed after each sampling (Figure 1D). Samples were collected at intervals of 4, 8 and 12 weeks in each season (Figure 1E).

### Sample collection, pre-processing and DNA extraction

During each sampling session, three monthly recruitment samples of each site were collected and then replaced with three new meshes. Three random samples from the seasonal succession frames were also collected. One new mesh sample was included for each site as a negative control. Sterile gloves and 10% bleach were used to minimise cross-contamination. The collected mesh samples were then stored in sterile 50 mL falcon tubes (Item No. 227-261; Cellstar^®^, Greiner Bio-One, Austria) placed on ice until transported to laboratory. Logged data were downloaded every sampling and loggers reset. In total n = 252, samples were obtained, including 216 biofouling samples and 36 field negative controls.

Wet weight (WW) data were recorded immediately after the samples arrived at the Plant & Food Research Institute (PFR) laboratory. The samples were then stored at −80°C for at least 24 hrs before further processing. Freeze drying (24-48 hrs at −57°C; Gamma 1-16 LSC, Christ) and bead beating (2 min, 1500 RPM; 1600 MiniG Spex SamplePrep) were used for sample homogenisation at the Cawthron Institute laboratory. The samples were then returned to the PFR laboratory for dry weight (DW) data recording. A subsample (100 mg) was taken from each 50 mL tube and transferred to a 5 mL PowerBead Pro tube (DNeasy PowerSoil Pro DNA Isolation Kit, QIAGEN, MOBIO, Carlsbad, USA) with DNA then extracted following the manufacturer’s protocol. A negative DNA extraction control sample was included in every extraction run. DNA quality and quantity control were performed on a Nanodrop photometer (Implen Nanophotometer, Munich, Germany). Each extracted DNA sample was stored in 50 µL elution buffer at −20°C until library preparation.

### Library preparation and high-throughput sequencing

The V4 region of the eukaryotic nuclear small-subunit ribosomal RNA (18S rRNA) gene was amplified by Polymerase Chain Reaction (PCR) using the primer set Uni18SF: 5’-AGG GCA AKY CTG GTG CCA GC-3’ and Uni 18SR: 5’ -GRC GGT ATC TRA TCG YCT T-3’ (Zhan et al. 2013), with the Illumina overhang adapters for dual indexing (Kozich et al. 2013). DNA samples were diluted to 1:25 (for samples with concentrations between 100 ng/µL and 250 ng/µL) and 1:50 (for samples with concentrations higher than 250 ng/µL) to improve the target gene amplification yield. PCR reactions were performed in a thermal cycler (Eppendorf^®^ Mastercycler, Hamburg, Germany) in a total volume of 50 µL, each reaction including 25 µL of 2× MyFi™ PCR Master Mix (Bioline Meridian Bioscience, Memphis, Tennessee, USA), 2 µL of each primer (10 µM stock), 2 µL of template DNA, and 19 µL of double-distilled water (ddH_2_O). The PCR cycles were 95 °C for 3 min followed by 37 cycles of 94 °C (20 s), 54 °C (20 s), and 72 °C (30 s) with a final extension at 72 °C for 7 min (Audrézet et al. 2022). A PCR negative control (no DNA) was included in each PCR run. Quality controls of the amplicons were performed on randomly selected samples (n=66) using gel electrophoresis. A subset (25 µL) of each amplicon was purified and normalised to a concentration of 1-2 ng/µL using SequalPrep normalisation plates (ThermoFisher Scientific, USA). The normalised amplicons were then sent to Sequench Ltd in Nelson, New Zealand, for NEXTERA indexing and paired-end sequencing (2 × 300 bp) using an Illumina Nextseq^TM^ platform. All raw sequence reads were deposited in the NCBI sequence read archive (SRA), under BioProject PRJNA1302565.

### Bioinformatics and statistical analyses

The Illumina overhang adapters were removed by the Nextseq^TM^ platform following sequencing. Raw sequencing data in the FASTQ files were subjected to demultiplexing and primer removal (with 19 bp overlap) using CUTADAPT, version 4.9 (Martin, 2011) in R. Quality filtering and denoising were followed by the parameters (truncLen=c(280,280), maxN=0, maxEE=c(2,2), truncQ=2) based on the read quality score using the R package DADA2, version 1.26.0 (Callahan et al. 2016). Forward and reverse reads were merged with a minimum 10 bp overlap with no allowed mismatches. Chimeras were removed using the default ‘consensus’ method. The filtered sequences were then taxonomically assigned to the SILVA reference database for the 18S rRNA gene, 138.1 SSU Ref NR 99 (Yilmaz et al. 2014). Potential contamination sequences found in the negative controls were removed using the R package microDecon, version 1.0.2 (McKnight et al. 2019). Sequencing depth and diversity coverage of all samples were inspected via rarefaction curves generated by the ‘ggrare’ function from the R package ranacapa, version 0.1.0 (Kandlikar et al., 2018).

Wet and dry weight data were visualised with line plots using the R package ggplot2, version 3.3.6 (Wilkinson 2011). Alpha-diversity metrics (observed richness and Shannon index) were calculated based on the rarefied data (3241 reads/sample) using the R package phyloseq, version 1.40.0 (McMurdie & Holmes 2013). The effects of exposure and season and their interactions on weight and alpha diversity were tested using two-way analysis of variance (ANOVA; Sthle & Wold 1989). Assumptions for normality and heterogeneity of variance were tested using Shapiro-Wilk and Levene’s tests (assumptions were met if p > 0.05). Tukey’s honestly significant difference (Tukey’s HSD) post-hoc test was performed for pairwise comparisons. If any assumptions were violated, aligned rank transform (ART) as a non-parametric approach was used for ANOVA using the R package ARTool, version 0.11.2 (Wobbrock et al. 2011). Where ART was applied, pairwise comparisons were conducted using the R package emmeans, version 1.11.2 (Lenth 2025).

Community composition was analysed on the unrarefied and normalised proportional data using phyloseq and visualised with stacked bar plots using ggplot2 and the R package khroma, version 1.16.0 for colour palette (Frerebeau 2025). The classes of the most abundant 50 species were selected for plotting as the most representative taxonomic groups. Non-native species analysis was performed based on the latest 161 species listed by Biosecurity New Zealand using the pest alert tool, following 99.4% of the minimum sequence identity match and 400 bp of the minimum sequence length (Zaiko et al. 2023).

Beta-diversity was analysed for the effects of environmental factors on community composition (Whittaker 1960). Two-way permutational analysis of variance (PERMANOVA) on rarefied data was used for testing the effects of the two factors (e.g., season and exposure, temperature and wind fetch) and their interactions. The function “adonis_pq” (permutations = 999, method = “bray”, by = “terms”) from the R package MiscMetabar (Taudière 2023) was used for phyloseq objects, based on the “adonis2” function from the R package vegan, version 2.7-1 (Oksanen et al. 2025). Before that, homogeneity was tested for the factors’ interactions with permutation of dispersions (PERMDISP) (assumption was met if p > 0.05) using the “betadisper” function from vegan. Analysis of similarities (ANOSIM) was used in vegan if the assumption was rejected (Clarke and Green 1988). Post-hoc pairwise comparisons were conducted using the R package pairwise Adonis, version 0.4 (Martinez 2020). The rarefied data were visualised using non-metric multidimensional scaling (NMDS) plot with Bray-Curtis dissimilarity distances across the two main factors (Kruskal 1964). Further than this, temperature and wind fetch as the representative factors of season and exposure were quantified and visualised using ggplot2 and the R package windfetch, version 2.1-1 (Seers 2018). The seasonal succession data were used for testing the effects of season and exposure, and the monthly recruitment data were used for testing the effects of temperature and wind fetch.

## Results

All amplicons tested by gel electrophoresis had clear target bands (around 600 bp). Close double bands were observed in some amplicons, a pattern commonly seen in samples with multiple species and amplified by universal primers with low specificity. No visible bands were found in negative controls. The raw sequencing yield of the 216 amplicons (negative controls excluded) were 32,486,494 sequence reads (average of 150,400 reads per amplicon). After data cleansing, the total number of reads was reduced to 21,309,064 (average of 98,653 per amplicon). The final 20,153,751 reads (average of 93,304 per amplicon) were retained after decontamination removal based on the negative controls, with a total of 10,153 amplicon sequence variants (ASVs). One amplicon with no reads was excluded for the downstream analysis. The full list of the read count of each sample at each cleansing step is provided in Table S1.

### Biomass and alpha diversity to measure seasonal accumulation patterns

The WW and DW accumulation patterns at the exposed sites were significantly different between summer and winter (p = 0.0264 WW; p < 0.0001 DW), whereas similar seasonal patterns were found from the sheltered sites (p > 0.5 for both WW and DW) (Figure 2A & 2B). For the exposed sites, biomass accumulated faster and more continuously in winter, with significantly higher quantities after 12 weeks than 4 weeks (p < 0.0001 for both WW and DW at all the exposed sites) (Figure 2A & 2B). In summer, dramatic weight increases and decreases were observed at the exposed sites (p < 0.001 for the WW and DW increases from week 4 to 8 and the WW decreases from week 8 to 12 at E1 and E3; p < 0.0001 for the WW and DW increases from week 8 to 12 at E2) (Figure 2A & 2B). During the first four weeks post-deployment, biomass accumulation was generally higher at sheltered sites compared to exposed sites in both seasons (p < 0.05), except the DW in summer (Figure 2B). Beyond this initial period, changes in biomass at sheltered sites were more gradual, whereas exposed sites exhibited greater temporal variability (Figure 2). Higher average biomasses (both WW and DW) were accumulated at the sheltered sites in summer (p < 0.05), whereas heavier average DW was found at the exposed sites in winter (p = 0.0152) (Figure 2A & 2B). Higher water content of the biomass was also observed from the exposed sites in summer compared to winter (Figure 2). The full list of the ANOVA and post-hoc pairwise comparison results of biomasses are available in Table S2.

**Figure 2.**
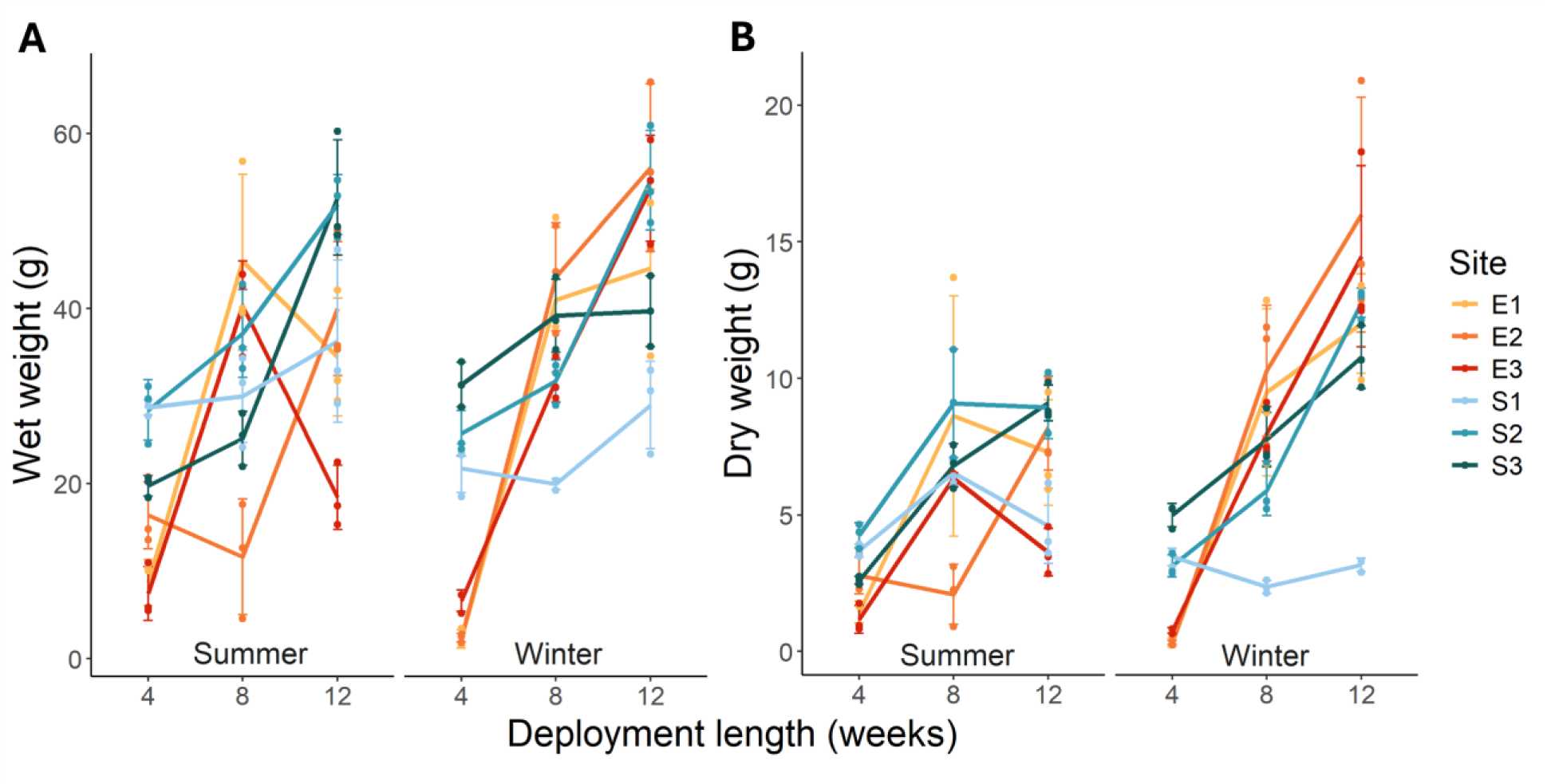
Biomass of the successional samples from each site in two seasons. Line graph with data points and error bars of wet weights (A) and dry weights (B) of the seasonal succession samples at the sites from two exposures (displayed in warm colours for the exposed sites and cold colours for the sheltered sites) in summer and winter (week 4 to 12), respectively.

The observed richness of sheltered sites was higher than exposed sites in both seasons (p < 0.05), especially in summer (p < 0.0001) (Figure 3A). No significant seasonal differences were found within the same exposure types (p > 0.05) (Figure 3A). Numerous temporal fluctuations were observed between both season and exposure (E1 & S1 in summer and S2 & E1-3 in winter) (Figure 3A). For the Shannon index, the growing patterns of the sheltered sites in both seasons and the exposed sites in summer were similar, except for a drop recorded at S1 at the 8-week growth point during summer (p = 0.0001) (Figure 3B). In winter, dramatic increases at two exposed sites (E1 & E3) occurred between weeks 4 and 12 (p < 0.0001), ranging from 0.38 to 2.64 (Figure 3B). The full list of the ANOVA and post-hoc pairwise comparison results of alpha diversity matrix is available in Table S3.

**Figure 3.**
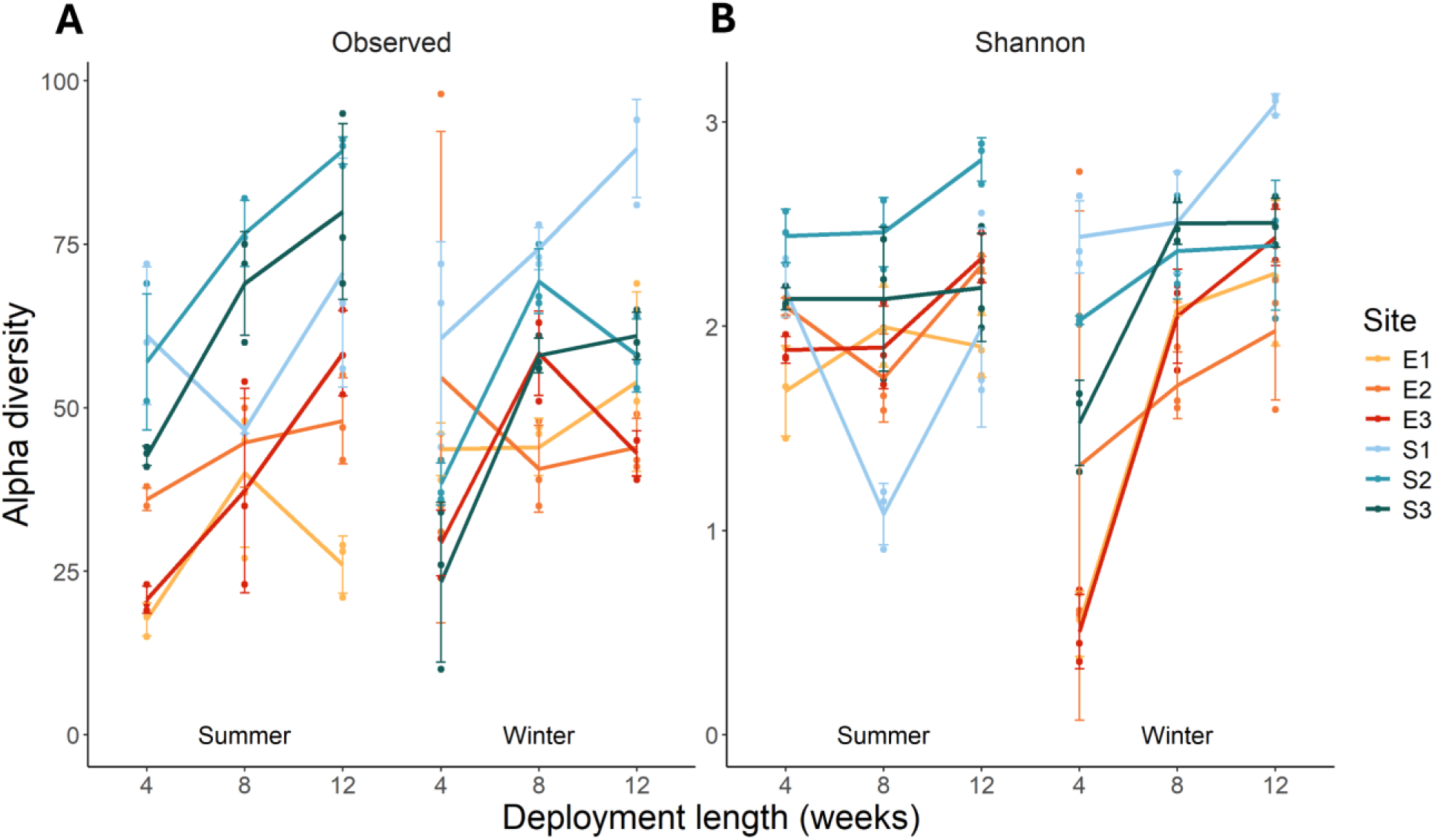
Alpha diversity of the successional samples from each site in two seasons. Line graph with data points and error bars of observed richness (A) and Shannon diversity (B) of the seasonal succession samples at the sites from two exposures (displayed in warm colours for the exposed sites and cold colours for the sheltered sites) in summer and winter (week 4 to 12), respectively

### Biofouling community composition analyses to investigate seasonal dynamics

The most abundant 50 ASVs were assigned to 13 classes and 22 species, with two classes accounting for more than 50% of the total number of reads (29.83% of Malacostraca and 36.38% of Hydrozoa) (Table S4). Strong seasonal patterns were found at the exposed sites, with Oligohymenophorea (26.85% of the total reads from the exposed sites in summer), Copepoda (17.01%, mainly including 13.55% of *Harpacticus* sp. and 3.13% of *Temora turbinata*) and Malacostraca (46.43%, including 35.78% of *Jassa slatteryi*, 5.25% of *Jassa falcata* and 5.40% of *Caprella equilibra*) dominated in summer, whereas Hydrozoa (76.72%, including 52.54% of *Coryne eximia* and 24.18% of *Obelia geniculata*) and Malacostraca (13.47%, mainly including 6.80% of *Jassa slatteryi* and 6.49% of *Cyclograpsus cinereus*) dominated in winter (Figure 4). A more obvious successional trend was observed at exposed sites in summer compared to winter. In summer, Gastropoda and Malacostraca decreased from weeks 4 and 8, while Copepoda increased from week 8 at all three sites (Figure 4). Comparatively, Hydrozoans were dominant throughout the winter season at all three exposed sites with only small increases in the amount of Malacostraca and other classes occurring from weeks 8 and 12 (Figure 4).

**Figure 4.**
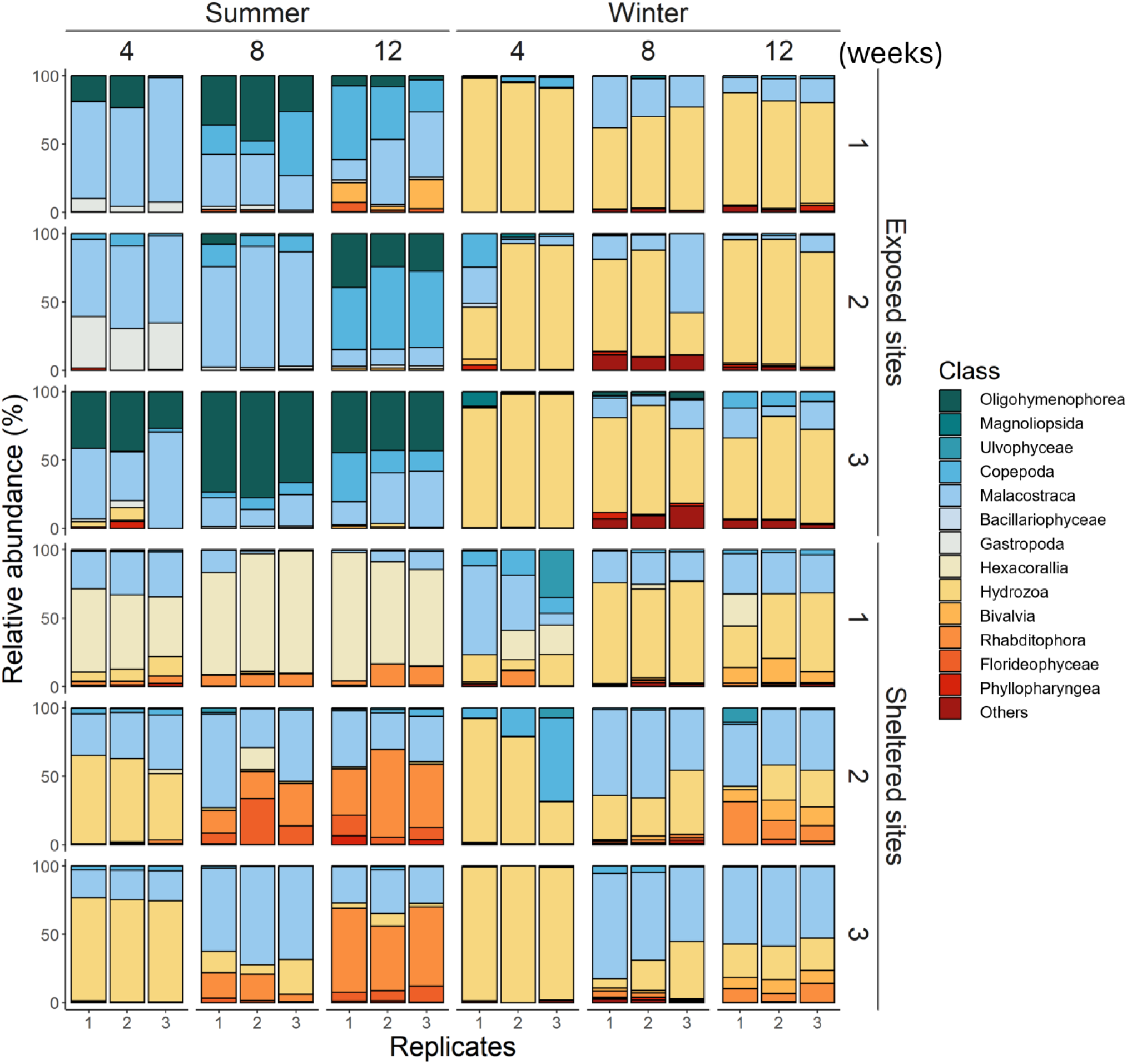
Class-level composition of eukaryotic biofouling communities from the successional samples across two locations (sheltered and exposed) and two seasons (summer and winter). Relative abundances (%) of the most abundant 50 ASVs (displayed at class taxonomy level in different colours) from the seasonal succession samples at the sites of the two exposures in summer and winter (week 4 to 12), respectively. The three replicates of each treatment were displayed together.

Although the seasonal patterns were less pronounced at the sheltered sites, distinct site differences were observed, especially in summer, with Hexacorallia (*Viatrix globulifera* at 69.22%) dominant at S1 (Figure 4). Relatively similar seasonal patterns were found at S2 and S3. Hydrozoa (32.09%-42.82%, including *Obelia geniculata* at 24.42%-38.49% and *Coryne eximia* at 4.33%-7.67% for both seasons) grew rapidly in the first 4 weeks, then reduced from week 8, and Malacostraca (32.45%-42.61%, including *Jassa slatteryi* at 20.76%-26.57% and *Caprella equilibra* at 11.69%-16.04% for both seasons) and Rhabditophora (22.81% and *Notoplana australis* at 5.36% in summer and winter, respectively) appeared from weeks 4 and 8 until week 12 in both seasons (Figure 4). In winter, more Hydrozoa species were detected at S1 than the other two sheltered sites from week 8, and both Ulvophyceae and Copepoda were observed during the first 4 weeks at S1 and S2 (Figure 4). The read counts and percentages of the classes and species from both seasons and locations are available in Table S4.

For non-native species, all detected ASVs could be assigned to nine species, with *Jassa slatteryi* (23.72% of the total non-native species reads), *Obelia geniculate* (34.84%) and *Coryne eximia* (41.44%) as the most abundant of these. Similar to the full community compositions above, strong seasonal patterns from the exposed sites were found in the non-native species, with 98.26% of the reads in summer were *Jassa slatteryi* and 93.28% of the reads in winter were Hydrozoan species (*Obelia geniculata* at 18.82% and *Coryne eximia* at 74.46%) (Figure 5). Seasonal variation in non-native species composition was less pronounced at sheltered sites. However, *Jassa slatteryi* was more prevalent in summer, particularly at S1 and S2, while *Obelia geniculata* was more common in winter. Additionally, *Cladophora ruchingeri* was detected in low abundance from week 8 in summer and throughout weeks 4 to 12 in winter at both S1 and S2 (Figure 5). *Coryne eximia* was more frequently observed at exposed sites, particularly during winter, whereas *Obelia geniculata* was more abundant at sheltered sites across both seasons (Figure 5). The read counts and percentages of the non-native species from both seasons and locations are available in Table S5.

**Figure 5.**
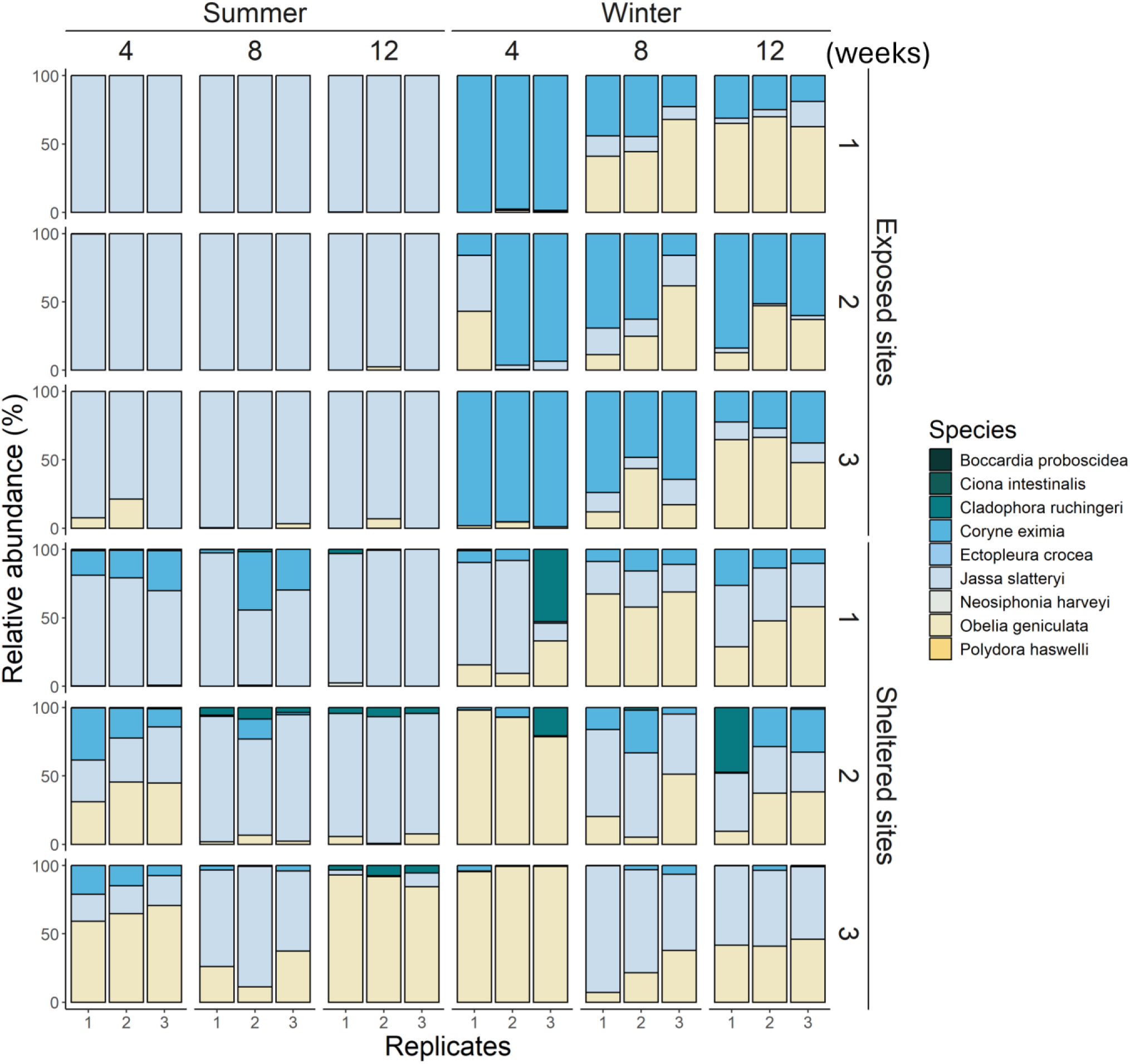
Species-level composition of the detected non-native species from the successional samples across two locations (sheltered and exposed) and two seasons (summer and winter). Relative abundances (%) of all the detected non-native species (displayed at species taxonomy level in different colours) from the seasonal succession samples at the sites of the two exposures in summer and winter (week 4 to 12), respectively. The three replicates of each treatment were displayed together.

### Beta diversity to access the impact of season and exposure on seasonal succession patterns

Both season and exposure, and their interaction had significant effects on the seasonal succession patterns of community composition (p = 0.001 for both factors and their interactions). The summer patterns from the exposed sites were significantly more isolated from the other treatments (p = 0.006) (Figure 6). The patterns of the exposed sites in winter and the sheltered sites in both seasons were closer to each other but still significantly different across all sites and all sampling points (p = 0.006) (Figure 6). Furthermore, significant seasonal succession trends were found in both seasons and at both exposures (p < 0.05), except for the communities between week 8 and 12 at sheltered sites (p = 0.414) and between week 4 and 8 at exposed sites (p = 0.144) in summer (Figure 6). Site differences within the same season and exposure were also observed, with the patterns at E3 being significantly different from E1 and E2 (p = 0.003), and S1 significantly different from the two other sheltered sites (p = 0.003) in summer (Figure 6). The full list of the ANOSIM, PERMANOVA and post-hoc pairwise comparison results of beta diversity matrix is presented in Table S6.

**Figure 6.**
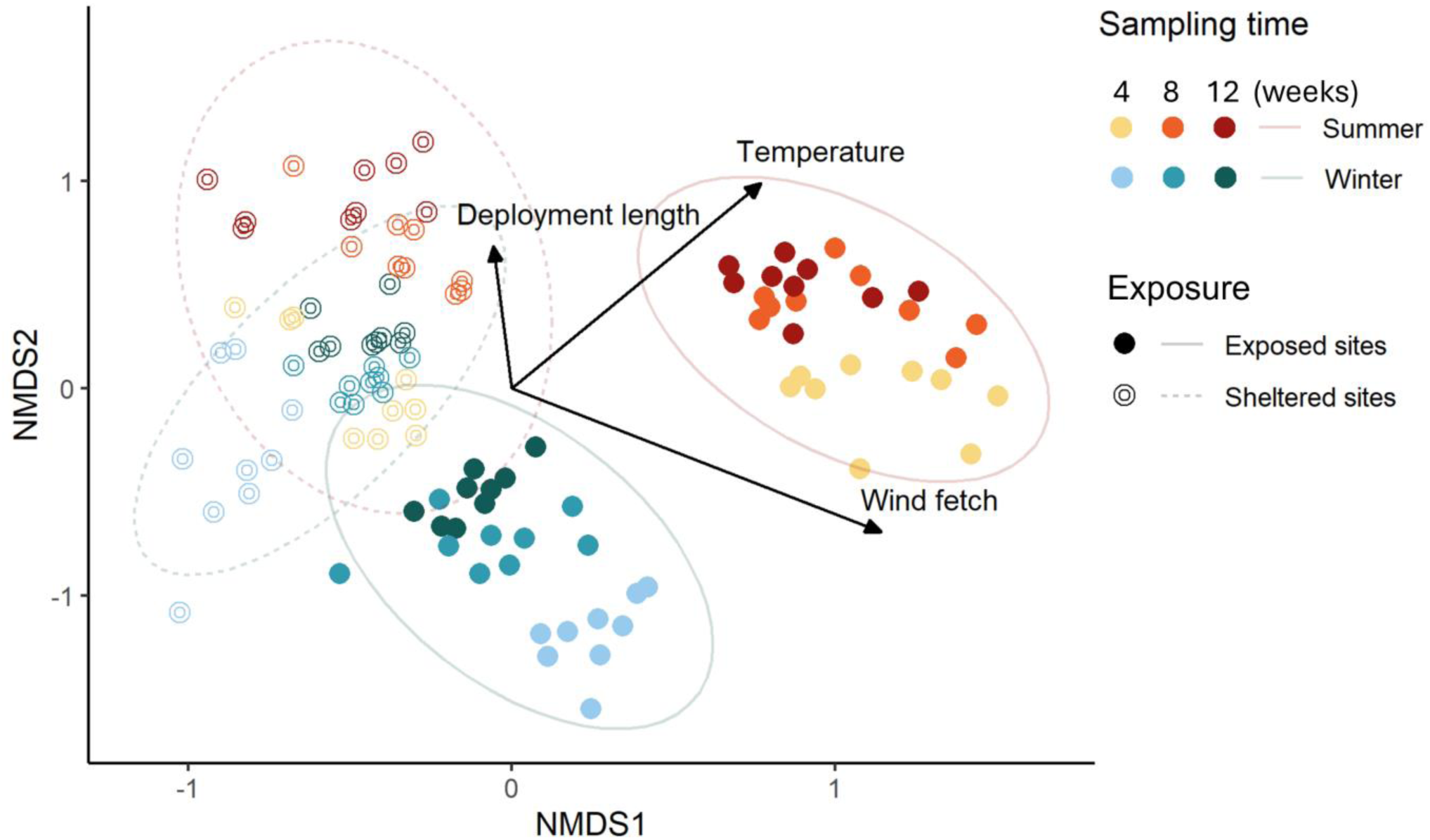
Beta diversity of the seasonal succession samples across two exposures and two seasons. NMDS plot of the seasonal succession samples at two exposures (displayed in different shapes) in summer and winter (week 4 to 12, displayed in warm colours for summer and cold colours for winter), respectively

### Effects of temperature and wind fetch on monthly recruitment patterns

The averaged wind fetches for each of the exposed sites (78,674.24 m at E1, 80,482.00 m at E2, 80,600.62 m at E3) were significantly higher (around 40 times, p < 0.0001) than the sheltered sites (2,535.90 m at S1, 2,450.51 m at S2, 3,336.66 m at S3) (Figure 7A & 7B). No significant wind fetch difference was found within the same exposure (p = 0.615) (Figure 7A & 7B). Summer temperatures were significantly higher than the temperatures in winter at both exposures (p < 0.0001) (Figure 7C). Summer temperatures at exposed sites (20.05 ± 0.63 °C) were significantly higher (p < 0.0001) than those at sheltered sites (18.31 ± 0.63 °C) (Figure 7C). There was no significant temperature difference (p = 0.1190) between the two exposure types in winter (13.17 °C ± 0.76 °C) (Figure 7C). Both temperature and wind fetch, and their interaction had significant effects on the monthly recruitment patterns (p = 0.001 for both factors and their interactions). The communities from both high- and low-temperature environments (summer and winter) at both wind fetch levels (exposed and sheltered) were significantly different from each other (p = 0.006). Some significant site differences within the same level of wind fetch in both seasons were found (p = 0.003 for S1 differing from S2 & S3 in summer and S3 in winter, p = 0.003 and 0.024 for E2 differing from E1 & E3, respectively), indicating site-specific variation under similar exposure conditions (Figure 8). From the wind fetch perspective, locations with both levels of wind fetch had distinct seasonality (p = 0.006), with larger seasonal variances for the location with significantly higher wind fetches (Figure 8). Different monthly community patterns at different sampling points from the same temperature and wind fetch levels were also observed (p < 0.01, except for the communities between week 4 and 8 at exposed sites in summer with p = 0.192, between week 8 and 12 at sheltered sites in summer with p = 0.516, and between week 8 and 12 at exposed sites in winter with p = 0.087) (Figure 8). The full lists of the temperature and wind fetch calculations, ANOSIM, PERMANOVA and post-hoc pairwise comparison results of beta diversity matrix can be found in Table S7.

**Figure 7.**
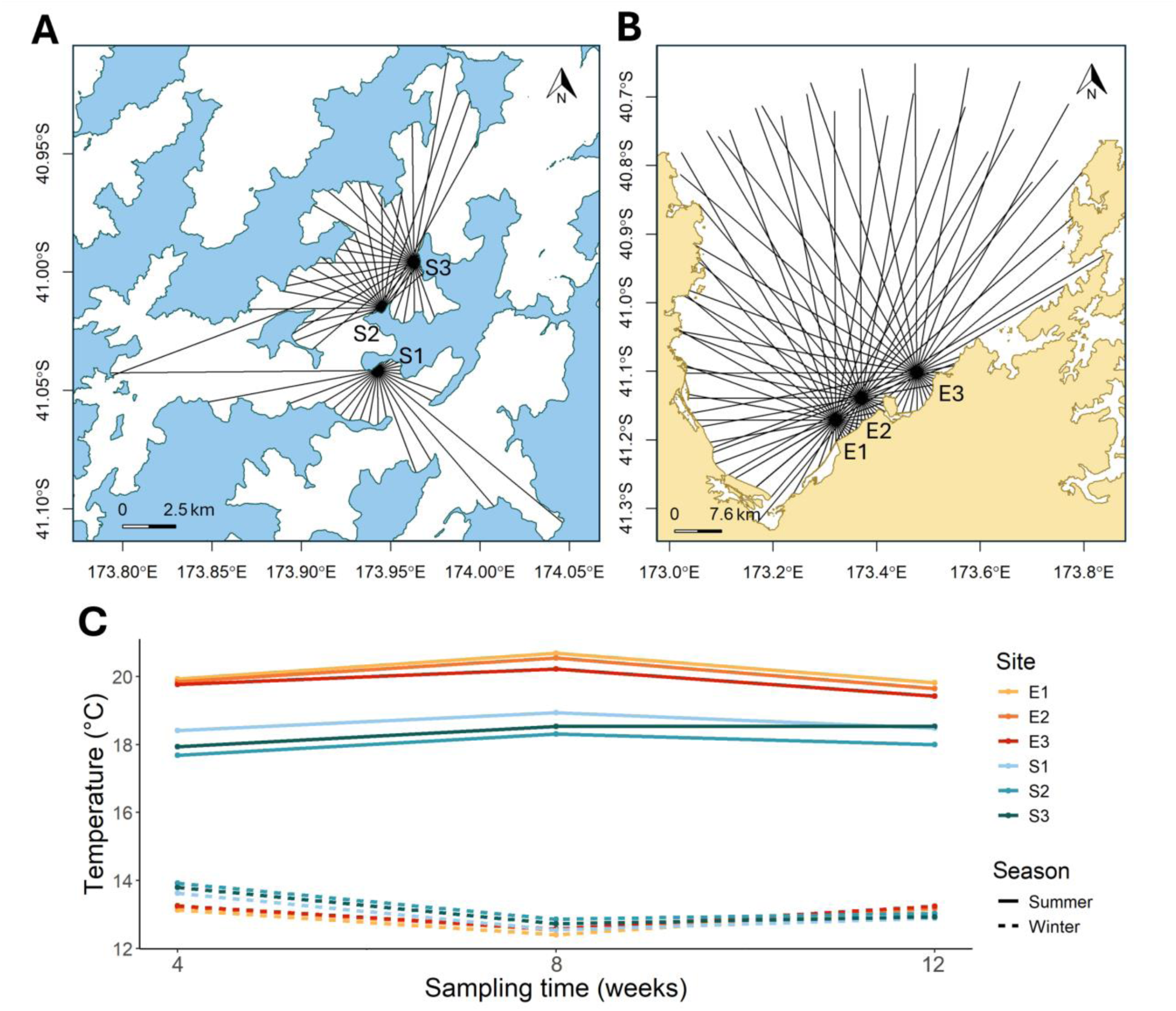
Wind fetch of each site and the temperature changes in each sampling season. (A) Visualised wind fetch (distances from the site to the coasts in all directions) of each sheltered site; (B) visualised wind fetch of each exposed site; (C) line graph of the average monthly water temperatures at each site (displayed in different colours) at each sampling point in summer and winter (displayed in different line types).

**Figure 8.**
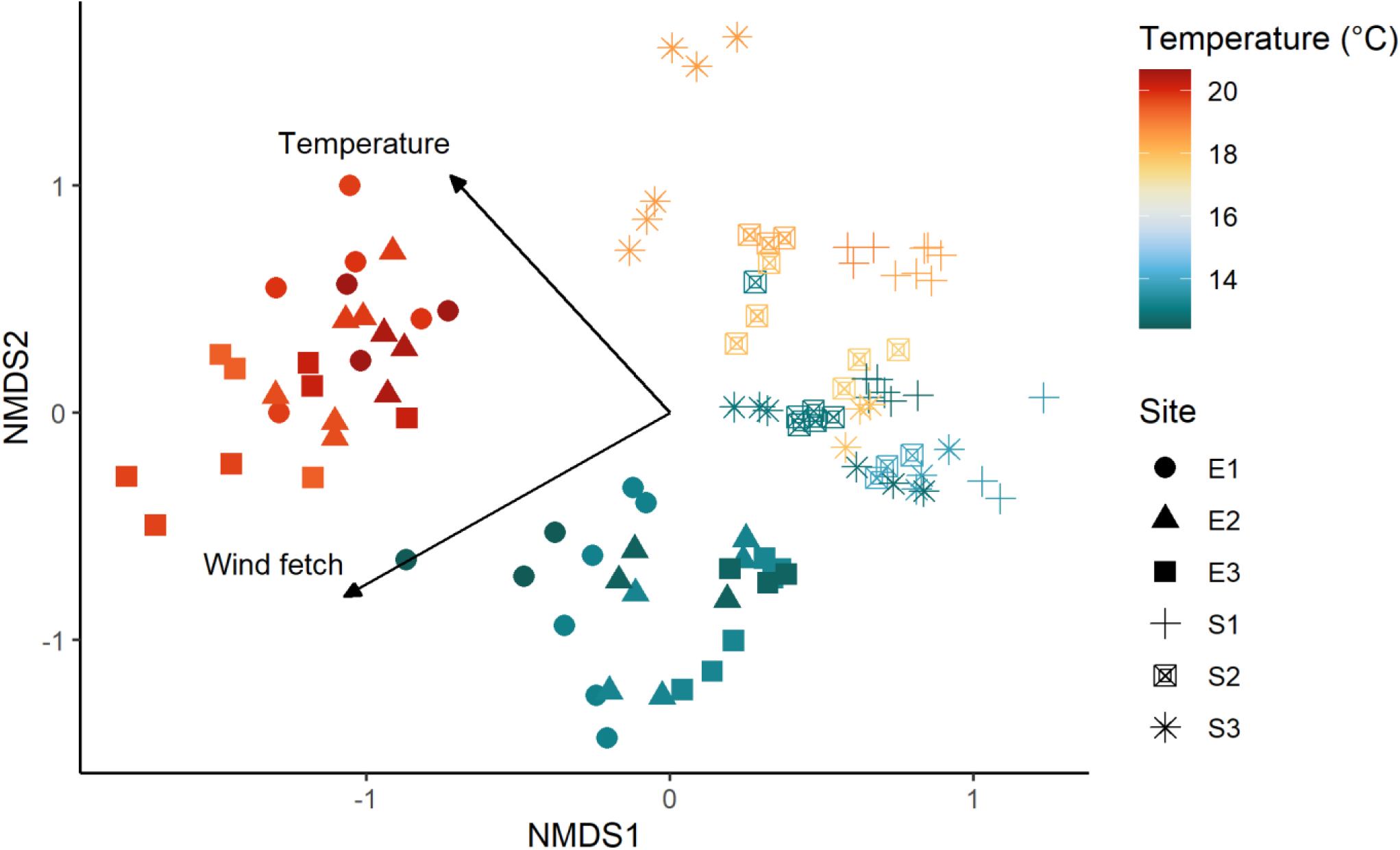
Beta diversity of the monthly recruitment samples across six sites and different temperatures. NMDS plot of the monthly recruitment samples at six sites (displayed in different shapes) with continuous temperature changes in summer and winter (displayed in gradient colours). The wind fetch of the exposed sites and the sheltered sites were significantly different and were considered as categorical data, which can be represented by the sites with two levels of exposures.

## Discussion

### Biomass and diversity

In this study, strong seasonality and spatial variations were identified in biomass and diversity of eukaryotic biofouling, with temperature and wind exposure observed as key factors driving shifts in community patterns (Garner and Litvaitis 2013; Lord 2017; Visch et al. 2020; Ulaski and Konar 2024). Higher water temperatures and less exposure are generally considered as the factors that can accelerate biomass growth (Atalah et al. 2016; Khosravi et al. 2019). However, we found higher biomass and faster community development in colder and more exposed locations, indicating potential underlying drivers of these trends. A biofilm study reported lower bacterial species richness in summer compared to winter, which may reduce the availability of settlement cues and subsequently lead to decreased recruitment of eukaryotic fouling organisms (Chan et al. 2003). A similar phenomenon was noticed through visual observation in this study, where larger quantities of biofilm formed at the exposed sites during winter, before the growth of eukaryotic fouling community. To explain the reduction in WW/DW over summer, it was noted that amphipod-dominated communities were not tightly bound to the mesh and could be readily dislodged by increases in water flow or disturbances by marine animals. This was not observed among the more deeply “rooted” hydroid-dominated communities in winter. In contrast, the sheltered sites had more stable seasonal patterns, with similar biomass and slightly higher genetic diversity than the exposed sites. These findings suggest that marine environments with greater seasonal variability, such as fluctuations in temperature and water movement, tend to exhibit more pronounced differences in biofouling patterns between seasons.

### Community composition

The majority of the heavy and dense biomass at exposed sites during winter was attributed to two hydroid species *Obelia geniculata* and *Coryne eximia*, whereas summer communities were dominated by two amphipod species *Jassa slatteryi* and *Jassa falcata*. Previous studies suggest that the biomass accumulation and reproductive activities of hydroid species typically increase with rising summer temperatures (Puce et al. 2003; Slobodov and Marfenin 2004; Park and Hwang 2012). The enhanced hydroid growth observed in winter in this study may be attributed to their strong adhesive capabilities, the early occurrence of biofilm, and the intensified water movement at exposed sites during this season, which could facilitate nutrient availability (Hadfield 2011; Di Camillo et al. 2017; Chiswell et al. 2021). These hydroid species add structural complexity to underwater substrates providing habitat and food for small invertebrates, such as crabs (*Cyclograpsus cinereus*), which were frequently and exclusively found at the exposed sites in winter in this study. Comparatively, the soft and tubular structures constructed by amphipods using debris from biological and non-biological materials do not provide the same degree of habitat complexity (Beermann and Franke 2012). Although, seasonal differences at sheltered sites were less pronounced than at exposed sites, a greater diversity was detected, including algae such as *Cladophora ruchingeri* and *Polysiphonia* sp. Also, strong site differences were detected at sheltered sites especially in summer. The dominant anemone *Viatrix globulifera* and caprellid *Caprella equilibra* were detected at S1 (all samples) and S3 (only the third monthly recruitment samples), respectively. The anemone *Viatrix globulifera* (currently considered as a synonym of *Bunodeopsis globulifera*) has been detected in the sheltered area from previous studies, but the drivers of its accumulation have been less studied (Fletcher et al. 2023). Detecting the presence of cnidarians is particularly important as several species have been implicated in causing negative outcomes for animal welfare and productivity on salmon farms (Fletcher et.al, 2023). Detections such as this illustrates how fine-scale analysis of spatial and temporal settlement patterns can potentially be utilised to refine management approaches and the timing of interventions such as cleaning. In terms of the caprellid amphipods, we found rapid and displacing growths occurred on both clean mesh and developed biofouling structures, indicating this might be driven by relevant feeding or competitive behaviours (Guerra-García and Tierno de Figueroa 2009).

### Non-native species and species that pose known risks to aquaculture

Interestingly, the three most dominant species detected in this study (*Jassa slatteryi*, *Obelia geniculata*, and *Coryne eximia*), were all non-native to New Zealand (Zaiko et al. 2023). These three species are commonly associated with biofouling communities globally but remain understudied in New Zealand, particularly regarding their potential impacts on marine aquaculture (Fernandez-Gonzalez and Sanchez-Jerez 2017; Navarrete et al. 2019; Beermann et al. 2020; Leclerc et al. 2020). Besides accumulating large biomass on aquaculture infrastructure, the hydroid species have been reported to negatively impact the quality of commercial kelp and salmon species by extensively growing on the surface of aquaculture species and spreading pathogens (Park and Hwang 2012; Kintner and Brierley 2019). The filamentous green algae (*Cladophora ruchingeri*) found at the sheltered sites was also considered as a problematic species for weight increasing and resource competition with native mussels (Cahill et al. 2022).

Apart from non-native species, there were also a number of potentially native harmful species detected. A turbellarian flatworm species (*Notoplana australis,* Rhabditophora) was mostly detected at the sheltered sites, particularly in summer. This species has been found in weakened oysters from previous research and could potentially cause risks for other mollusc species (Lester 1989). There was no record showing any issues on aquaculture about the dominant turtle grass anemone species (*Viatrix globulifera*) detected at S1 but can cause nerve damage to humans due to its cnidocytes (stinging cells) (Burke 2002).

### Implications for industry management

A detailed understanding of biofouling growth patterns under varying environmental conditions, along with the identification of species with potential risks, can inform the development of more effective antifouling strategies and operational guidance (Bannister et al. 2019). This is particularly important for biofouling management in marine industries, where early prediction and prevention are generally more effective and preferred over post-treatment measures (Fitridge et al. 2012). Furthermore, the variability of biofouling settlement under different conditions is often underestimated, highlighting the needs of finer-scale, long-term monitoring using cost-effective and efficient monitoring tools. Based on the season and exposure-driven biofouling patterns found in this study, we suggest that marine structures such as aquaculture farms selected with high wind exposures are more likely to face stronger environmental variations and larger seasonal differences in biofouling community composition and growth rates, indicating tailored biofouling management methods to optimise operation efficiency and cost (Portas et al. 2023). For instance, using smoother-surfaced, antibiotic materials and performing more frequent cleaning could prevent the formation of biofilms and dense hydroid settlement during winter season, whereas less frequent general physical removal may be sufficient for the amphipod dominant communities in summer. In sheltered locations with less seasonal variation, applying consistent fouling management approaches throughout the year may be more effective, but further antifouling tools are recommended for greater diversity and higher risk of harmful fouling organisms. Factors that are not strongly or directly relevant to season and exposure, such as anthropogenic influences and larval supply distribution, need to be explored to explain the community variations within the same season and exposure (Pineda et al. 2007).

### Methodological considerations and future work

Metabarcoding enabled high-resolution detection of eukaryotic biofouling taxa, offering greater taxonomic coverage and efficiency compared to traditional morphology-based methods (Taberlet et al., 2012). However, limitations remain, particularly with the use of 18S rRNA universal primers, which can lack taxonomic resolution for some species, leading to potential false positives, false negatives, and increased sensitivity to cross-contamination (Hadziavdic et al. 2014). For instance, the sequence detected as the Chilean shore crab species *Cyclograpsus cinereus* at the exposed sites may be a closely related smooth shore crab species *Cyclograpsus lavauxi* native to New Zealand and the only known Cyclograpsus species distributed in the South Island. This is because there are currently no 18S rRNA gene sequences for *Cyclograpsus lavauxi* in publicly available reference databases (only one 16S rRNA mitochondrial gene sequence available under GenBank accession number AB440191.1), which means any 18S rRNA sequence for this species could match sequences from other *Cyclograpsus* species.

Notably, some previously reported pest species such as *Mytilus galloprovincialis* and *Undaria pinnatifida* were not detected at the sheltered sites (Watts 2014; Atalah et al. 2020), which may reflect primer bias, low read abundance, or incomplete reference databases. Additionally, while metabarcoding is powerful for presence and absence and diversity assessments, it is less reliable for quantifying biomass or absolute abundance due to potential biases introduced during library preparation (Lamb et al. 2019). Although we included WW and DW measurements to estimate biomass in this study, such physical quantification is challenging to apply on operational marine structures, underscoring the need for complementary, scalable quantification tools.

For future biofouling monitoring research in marine environment, integrating metabarcoding with complementary quantification and validation tools, such as standardised and automated image analysis, is likely to become a trend for rapid and effective community assessment for industry (Gormley et al. 2018; Casey et al. 2021). The cost of biofouling management could be substantially reduced if community monitoring can facilitate preventative application of antifouling technologies, and/or improves the timing and efficacy of post-treatment tools such as physical removal (Bannister et al. 2019). Further monitoring on other critical biofouling communities, such as diatoms and bacteria, could potentially explain the eukaryotic biofouling patterns in this study. Apart from prevention and mitigation, biofouling biomass can also be utilized by extracting its chemical and nutrient components and processing them into value-added products, such as aquafeeds or medical materials. Alternatively, biofouling could potentially be used to facilitate contamination or nutrient remediation in restorative aquaculture and provide income to offset management costs (Zettler et al. 2013; Blunt et al. 2018). Biofouling management is not a simple process of ‘organism removal’, and there is no ‘one size fits all’ solution for the global marine industries. Only when we implement targeted and comprehensive strategies can we ensure quality and sustainable marine industries while maintaining a healthy aquatic ecosystem.

## Conclusion

This study shows that seasonal variation and wind exposure significantly influence biofouling community structure and biomass on mesh substrate. Using mesh material provided a realistic insight into fouling dynamics on industry-standard substrates. Exposed sites exhibited strong seasonal shifts, with dense hydroid assemblages in winter and more diffuse, amphipod-dominated communities in summer. Sheltered sites showed more stable biomass and higher species richness, with notable site-specific variability suggesting additional local drivers. These results underscore the need for context-specific antifouling strategies that consider both localised environmental conditions and material type. DNA metabarcoding proved effective for characterising eukaryotic biofouling communities, although limitations in quantification and primer specificity highlight the value of integrating complementary methods. Overall, this research informs targeted management approaches for marine industries and supports more sustainable biofouling mitigation practices.

## Acknowledgements

This research was funded by PFR’s SSIF Growing Futures investment. We also thank Ulla von Ammon and Michelle Scriver from Cawthron Institute for the valuable discussions for experimental design and bioinformatic analyses. Igor Ruza, Belinda Timms, Benie Chambers and Glen Aspin from PFR for the helps in field trips and lab work, and Sequench Ltd. for sequencing advice.

